# 2-APB arrests human keratinocyte proliferation and inhibits cutaneous squamous cell carcinoma *in vitro*

**DOI:** 10.1101/249821

**Authors:** Aislyn M. Nelson, Yalda Moayedi, Sophie A. Greenberg, Marlon E. Ruiz, Uffe B. Jensen, David M. Owens, Ellen A. Lumpkin

**Affiliations:** Department of Dermatology, Columbia University, College of Physicians and Surgeons, New York, New York 10032; Medical Scientist Training Program, Baylor College of Medicine, Houston, Texas 77030; Department of Physiology and Cellular Biophysics, Columbia University, College of Physicians and Surgeons, New York, New York 10032; Institute of Clinical Medicine, Aarhus University and Department of Clinical Genetics, Aarhus University Hospital, DK8200 Aarhus N, Denmark; Department of Pathology and Cell Biology, Columbia University, College of Physicians and Surgeons, New York, New York 10032

**Keywords:** keratinocyte differentiation, calcium channel

## Abstract

**Background:** The epidermis is a stratified epithelium whose differentiation program is triggered in part by calcium. Dysregulation of keratinocyte differentiation may lead to non-melanoma skin cancers, including cutaneous squamous cell carcinoma (cSCC). The compound 2-aminoethoxydiphenyl borate (2-APB) modulates calcium signaling by altering activity of calcium-permeable channels of the transient receptor potential (TRP) and ORAI families, and is therefore poised to govern signaling pathways that control the balance of keratinocyte proliferation and differentiation.

**Objective:** We sought to determine whether 2-APB alters differentiation of normal human keratinocytes and progression of human cSCCs models *in vitro*.

**Methods:** Primary human keratinocyte cultures were treated with 2-APB and levels of proliferation (EdU incorporation) and differentiation markers [quantitative PCR (qPCR)] were assessed. Human cSCC biopsies and cell lines were analyzed for TRP and ORAI gene expression via qPCR. cSCC cell lines were cultured in organtypic cultures and analyzed for growth and invasiveness after 2-APB or vehicle treatment.

**Results:** Culturing human keratinocytes with 2-APB arrested cell proliferation, triggered differentiation-gene expression and altered epidermal stratification, indicating that 2-APB application is sufficient to promote differentiation. In human organotypic cSCC cultures, 2-APB attenuated tumor growth and invasiveness. Finally, expression of a panel of 2-APB-targeted ion channels (TRPV3, TRPV1, TRPC1, OraI1, OraI2 and OraI3) was dysregulated in high-risk cSCC biopsies.

**Conclusions:** Collectively, these findings identify 2-APB as a potential therapeutic for high-risk cSCCs.

## Introduction

The incidence of human non-melanoma skin cancer is at an all-time high [1]. Although surgical treatment is adequate for low-risk cases, a subset of patients are at high risk for recurrence or metastasis [2]. For example, cutaneous squamous cell carcinomas (cSCCs) that develop in patients with chronic immunosuppression, such as in organ transplant recipients and patients with HIV, or tumors that develop in chronically injured skin, are frequently highly aggressive and potentially fatal [2]. Currently, there are no effective treatments to target tumors that develop in high-risk patients; therefore, new therapeutics are needed for non-melanoma skin cancers. Environmental factors, viral infections and genetic alterations that contribute to keratinocyte-derived skin cancers have been identified; however, a better understanding of normal keratinocyte physiology is critical for unmasking pathophysiological changes resulting in tumorigenesis.

In the healthy epidermis, calcium induces keratinocyte differentiation; therefore, Ca^2+^-permeable cation channels are poised to regulate this complex process. *In vivo*, the epidermis exhibits a calcium gradient necessary for normal structure and function: low calcium levels are found in the proliferative basal layer, whereas higher calcium levels induce keratinocyte differentiation in upper strata [3]. These conditions can be mimicked *in vitro* through a calcium-switch procedure. Normal keratinocytes proliferate in media containing 0.06 mM calcium and then are induced to differentiate by culturing them for a few days in >1 mM extracellular calcium. Calcium-triggered differentiation programs are dysregulated in non-melanoma skin cancers, leading to uncontrolled cellular proliferation. This suggests that calcium modulation is a potential therapeutic target for non-melanoma skin cancers.

Calcium concentrations are regulated in epidermal keratinocytes through store-operated calcium channels and non-selective ion channels on the plasma membrane, particularly those of the transient receptor potential (TRP) family. Store operated calcium entry (SOCE) occurs when stromal interaction molecules (STIM) 1 and 2 sense reduced endoplasmic reticulum calcium concentrations. In response, STIM 1 translocates to the membrane and stimulates calcium entry through ORAI channels. ORAI channels are thought to be primary components of the calcium release activated channel (CRAC) that mediates SOCE. In addition to ORAIs, there is some evidence for TRPC1 and TRPC4 participating in SOCE [4, 5]). Knockdown of both of these channels has been found to decrease SOCE currents. Upon calcium-induced keratinocyte differentiation, inositol triphosphate (IP3) induces release of ER calcium stores, which in turn induces CRAC currents [6]. The STIM/ORAI pathway has been implicated in tumor progression and metastasis in multiple models, including breast cancer and melanoma [7]. This observation, in combination with the role of ORAIs in regulating calcium homeostasis, has garnered increasing interest in ORAI modulation as a therapeutic strategy [7].

Several calcium-permeable channels of the TRP family have been implicated in pathological and normal keratinocyte function. Keratinocytes express TRPV3 and TRPV4 and pathological variants in these receptors cause epidermal abnormalities such as barrier defects and hyperkeratosis [8-11]. In humans, TRPV4 is strongly expressed in skin and downregulated in non-melanoma skin cancers [12]. In mice, TRPV3 regulates terminal keratinocyte differentiation via activation of epidermal growth factor receptor (EFGR) signaling [11]. Several reports indicate that keratinocytes also express TRPV1, TRPA1 and TRPC1 [14]. TRPV1 and TRPC1 are coupled to the EGFR pathway by means of cytokine factor release and calcium signaling, respectively [15, 16]. Increased expression and activation of EGFR correlates with poor prognosis in cSCC [17]. A number of EGFR inhibitors are FDA-approved for cSCC treatment but their efficacies are inconsistent and some are associated with severe skin toxicity [18]. Thus, in addition to being able to influence calcium influx, this link between TRP channels and the EGFR pathway indicates that these ion channels might regulate diseases associated with keratinocyte maturation.

To identify targeted therapies for cSCC, we sought to determine whether the calcium channel modulator 2-aminoethoxydiphenyl borate (2-APB) alters human keratinocyte behavior and progression of cSCCs *in vitro*. We found that 2-APB arrests human keratinocyte proliferation and promotes differentiation. In a pre-clinical *in vitro* model of human cSCC, 2-APB significantly reduced tumor growth and dermal invasion.

## Materials and Methods

Human tissue use was reviewed and exempted by the Columbia University Institutional Review Board.

### Cell culture

Normal human keratinocytes were isolated from human foreskins by two enzymatic dissociation steps, first in dispase then in trypsin. Cells were cultured in EpiLife (with human keratinocyte growth supplement; Invitrogen) for <5 passages. For all experiments with undifferentiated cells, keratinocytes were harvested at ~75% confluency. Differentiation was induced by adding CaCl_2_ to EpiLife media to reach a final [Ca^2+^] of 1.2 mM. Cells were typically cultured for 3 days under differentiation conditions before they were assayed.

Organotypic human skin and primary cSCC rafts were derived from normal human foreskin and cSCC keratinocytes, respectively, and propagated in keratinocyte growth medium as previously described [19]. In brief, human fibroblasts were seeded in a collagen matrix on a 70-μm filter insert and incubated at 37°C for 5–7 d. Human keratinocytes or SCC cells were then seeded on the surface of the fibroblast-collagen matrix and cultured for 2 days submerged in media to allow monolayer formation. After which, rafts were raised to the air-liquid interphase, exposing the keratinocytes or cSCC cells to air. The cultures were maintained for up to 14 d with media to the fibroblasts below changed every 2-3 d. At the time of raising, 50 μM 2-APB (Tocris) or vehicle (1% EtOH) was added the media.

### Histology

Sections were fixed in 4% paraformaldehyde, paraffin embedded and sectioned at 8 μm. Hematoxylin and eosin (H&E) stained sections were imaged with a brightfield microscope equipped with a 10X (0.3 NA) and 20X (0.4 NA) objectives and an Axiocam color CCD camera (Zeiss AxioObserver.Z1). Epidermal thickness was assessed by measuring nucleated layers with ImageJ software (NIH ImageJ, http://imagej.nih.gov/ij). The number of invading cells was determined by counting the number of round nuclei below the basement membrane per 10X field (fibroblast nuclei are flatten and were excluded). Four non-serial paraffin sections per experiment were examined. Six random frames per section were quantified.

### Proliferation

Cells were treated with 2-APB for 24 h at 37°C, exposed to EdU for 45 min, fixed and stained with DAPI for cell counting. EdU Click-It assays (Invitrogen) were used to assess cell-cycle entry. Three fields per well were assayed via a 20X objective and there were 4 wells per treatment in each experiment. The total number of Edu- and DAPI-positive cells were counted in each field of view and utilized to calculate the percentage of proliferating (Edu-positive/DAPI-positive) cells.

### qPCR

Complementary DNA was synthesized with oligo-dT primers and SuperScript III (Invitrogen). Primers were generated in Primer3. Primer pairs were optimized for qPCR and validated in control specimens. RNA was isolated using RNeasy Kit (Qiagen) and reversed transcribed. Four technical replicates were run for each primer set and cDNA sample. Standardized SYBR green amplification protocols were used on the StepOnePlus ABI machine as suggested by the distributor (Applied Biosystems). Melting curves were generated for all products to confirm a single amplicon for each product. To determine gene expression in each sample, cycle thresholds (C_T_) of the gene of interest were normalized to the reference gene GAPDH (ΔC_T_). Fold change was determined using the ΔΔC_T_ method where vehicle or normal keratinocytes were used as the calibrator (ΔΔC_T_=[(C_T_ (target gene)–C_T_ (reference gene)] – [C_T_ (calibrator)–C_T_ (reference gene)]. All gene amplifications were performed in quadruplicate.

### Statistics

Experimental replicates were performed using normal human epidermal keratinocytes isolated independently from human neonatal foreskin specimens (n=14 specimens). For each independent experiment, 3–8 technical replicates were performed. We obtained 12 individual cSCC and two normal skin biopsies. Four technical replicates were performed on each biopsy. Organotypic cultures were carried out in triplicate. Data are expressed as mean±SEM unless noted. Statistical significance was assessed with unpaired two-tailed Student’s *t* tests or one-way ANOVA with Bonferroni post *hoc* analysis (GraphPad Prism). For qPCR data, fold change in expression was log transformed and analyzed with one-way ANOVA with a Bonferroni *post hoc* analysis.

## Results

### 2-APB halts proliferation in human keratinocytes

At micromolar concentrations, 2-APB activates calcium-permeable TRPV channels, several of which have been showed to be expressed in human keratinocytes. As Ca^2+^ triggers the commitment switch from proliferation to differentiation in normal keratinocytes, we reasoned that increasing intracellular Ca^2+^ by incubating keratinocytes in 2-APB might be sufficient to induce this cell-fate switch in low-calcium growth media. To test this hypothesis, we cultured normal human keratinocytes for 24 h under low-Ca^2+^ conditions in the presence of 2-APB (0–100 µM).

First, we assessed cellular morphology, cell counts and cell cycle entry as evidenced by EdU incorporation. At 100 μM, 2-APB induced necrotic morphological changes and a loss of cell numbers, consistent with a previous report that saturating 2-APB concentrations induce keratinocyte cytotoxity (Fig. 1A-B) [21]. By contrast, at 50 μM 2-APB, keratinocytes appeared slightly larger than in vehicle-treated controls, which is a hallmark of keratinocyte differentiation (Fig. 1A) [20]. Moreover, there was a slight reduction in cell number and a complete inhibition of EdU incorporation, indicating cell-cycle arrest (Figs. 1B–C). Thus, we conclude that 50 μM 2-APB arrests proliferation without inducing necrotic changes in normal human keratinocytes. As proliferation arrest is a hallmark of cell-fate commitment, this observation is consistent with the hypothesis that modulating calcium channel activity can induce the fate switch from proliferation to differentiation in human keratinocytes.

**Fig 1.**
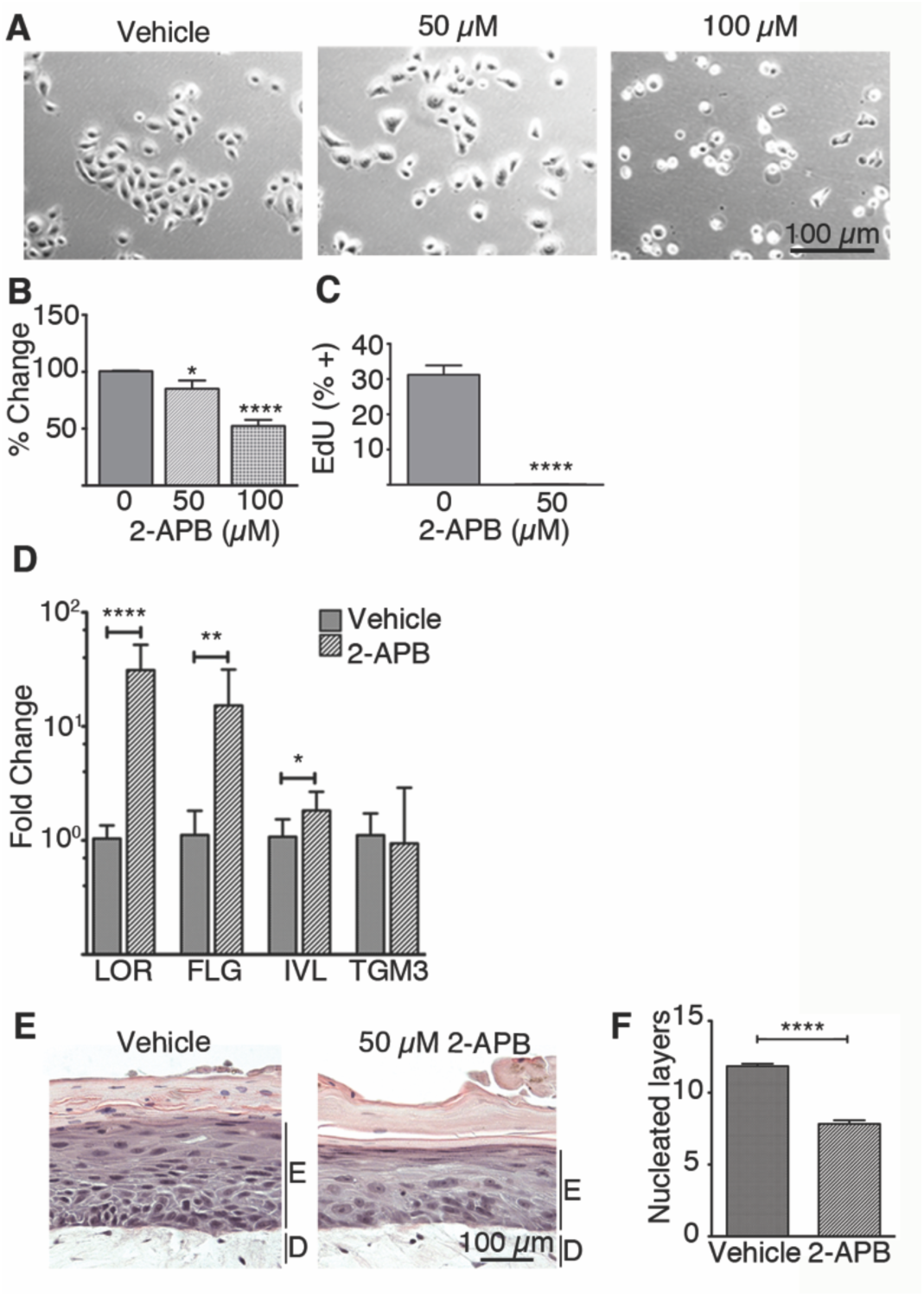
Constitutive treatment with 2-APB arrests human keratinocyte proliferation and promotes differentiation. **A-B.** Keratinocytes were cultured in a range of 2-APB concentrations. Phase contrast images correspond to concentrations plotted in B. **B.** Proportion of DAPI-labeled keratinocytes compared with vehicle-treated control cells (N=8 experimental replicates, *P<0.05, ****P≤0.0001, Student’s *t* test). **C.** Proliferation assays (45-min EdU pulse) after 24-h treatment with vehicle or 50 μM 2-APB (N=5 independent experiments, ****P≤0.0001, Student’s two-tail *t* test) **D.** Gene expression levels compared by qPCR (Loricrin: LOR, Filaggrin: FLG, Involucrin: IVL, Transglutaminase 3: TGM3). Fold increase in transcript levels in keratinocytes treated with 50 μM 2-APB compared with vehicle treatment is plotted (N=3 independent experiments; *P<0.05, **P≤0.01, ****P<0.0001, Student’s *t* test). **E.** Organotypic 3D human skin equivalents treated with vehicle (left) or 50 μM 2-APB (right). F. Mean number of nucleated epidermal cell layers is plotted (n=26–28 sections from two rafts per treatment, E=epidermal layer, D=dermis, ****P<0.0001). Plots depict Means±SEM.

We next asked whether incubation with 50 µM 2-APB promoted expression of well-established keratinocyte differentiation genes [22]. In low-Ca^2+^ growth media, loricrin (LOR) and filaggrin (FLG) transcript levels were induced more than 15-fold in keratinocytes treated for 24 h with 50 µM 2-APB compared with vehicle controls (Fig. 1D). By comparison, we observed limited change in expression of involucrin (IVL) and transglutaminase 3 (TGM3; Fig. 1D). This might be due to the 24-h 2-APB incubation period, as these late differentiation markers are typically not expressed in culture until 48 h after the induction of differentiation [23]. Alternatively, 2-APB might preferentially regulate early differentiation genes. These data indicate that constitutive treatment with 50 µM 2-APB in low-Ca^2+^ media is sufficient to commit human keratinocytes to a differentiated state.

Since 50 µM 2-APB induced proliferation arrest and promoted keratinocyte differentiation *in vitro*, we reasoned that 2-APB might alter epidermal stratification. We tested this prediction with human organotypic 3D skin equivalent models [24, 25]. Human keratinocytes were seeded on dermal matrices, allowed to form a monolayer, raised to an air-liquid interface to induce stratification and then treated for 7 d with either vehicle or 50 µM 2-APB (Fig. 1E). 2-APB-treated skin equivalents displayed a one-third reduction in nucleated epidermal layers (Fig. 1F). These data extend observations in two-dimensional keratinocyte cultures by demonstrating that constitutive exposure to 2-APB alters human keratinocyte behavior in a stratifying epidermis.

### Expression of 2-APB targets is dysregulated in human cSCC

To determine whether expression of calcium channel genes is dysregulated in SCC, we quantified expression of a subset of TRP- and ORAI-channel genes that are modulated by 2-APB (Table 1) and that are expressed in keratinocytes. A panel of human high-risk cSCC specimens was analyzed. We found that TRPV1, TRPV3, and TRPC1 gene expression levels were significantly dysregulated in >60% of cSCC biopsies compared with normal human skin (Fig. 2A-C). Of note, TRPV3 was significantly different from control skin in all but one tumor tested (Fig. 2C). ORAI1, ORAI2, and ORAI3 were significantly dysregulated in 33-50% of tumors tested. By comparison, expression of signature SCC biomarkers Cyclin D1 (CCND1) and EGFR [17] was altered in 33% and 41% of patient specimens, respectively (Fig. 2G-H). Although a more extensive survey is needed determine whether 2-APB targets might serve as biomarkers for high-risk cSCCs, these results illustrate that TRP and ORAI channel expression is dysregulated in a majority of human cSCCs examined.

**Fig. 2.**
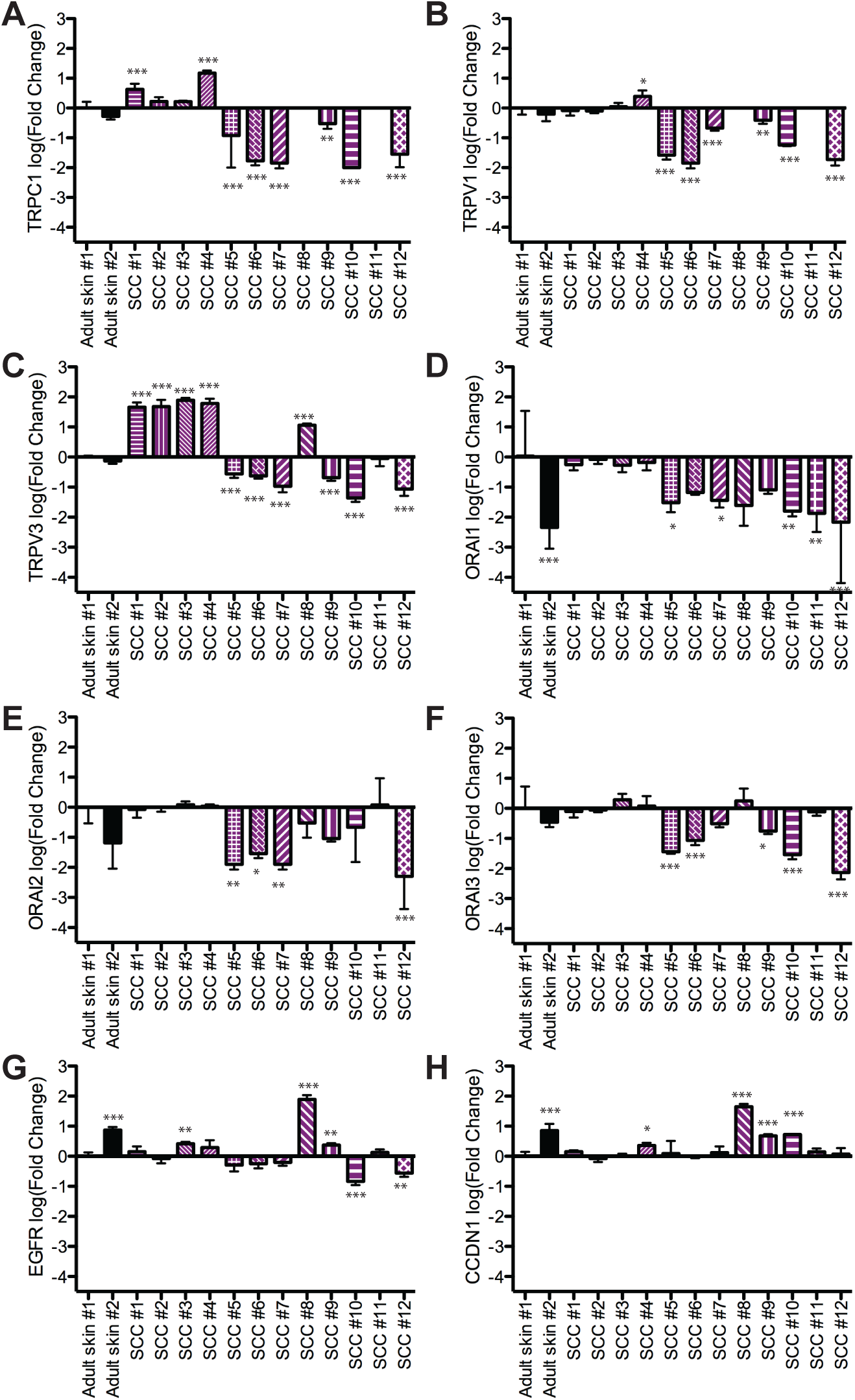
2-APB target channels are dysregulated in human cSCC. Gene expression levels of ion channels targeted by 2-APB assessed by qPCR in high-risk cSCC biopsies (purple patterns) and normal adult skin (black). **A.** TRPC1 **B.** TRPV1 **C.** TRPV3 **D.** ORAI1 **E.** ORAI2 **F.** ORAI3. For comparison, levels of EGFR (**G**) and CCND1 (**H**) are shown. Fold increase compared with normal adult skin sample #1 is plotted (N=4 technical replicates per sample, *P<0.05, **P<0.01, ***P<0.001, ****<0.0001 two-way ANOVA with Bonferroni *post hoc*). Plots depict Means±SDs.

**Table 1.** Ca^2+^ Channels modulated by 2-APB.

| **Channel** | **Action** | **Range (μM)** | **Reference** |
| --- | --- | --- | --- |
| IP3R | Inhibition | 10-100 | [42] |
| ORAI1 | Inhibition | 5-50 | [35, 36] |
| ORAI2 | Inhibition | 50 | [43] |
| ORAI3 | Activation | 1-100 | [36, 44] |
| TRPC1 | Activation | 5-10 | [45] |
| TRPC1 | Inhibition | 50-100 | [45] |
| TRPC3 | IP3-dependent inhibition | 30-75 | [46, 47] |
| TRPC6 | Inhibition | 10-300 | [48] |
| TRPM8 | Inhibition | 3-100 | [48] |
| TRPV1 | Activation | 30-1000 | [48] |
| TRPV2 | Activation | 50-1000 | [48] |
| TRPV3 | Activation | 3-1000 | [34, 48] |

### 2-APB reduces cSCC tumor growth and invasion in human preclinical models

To assess the functional effects of 2-APB in human cSCC keratinocytes, we employed two cell lines derived from human cSCC tumors (SCC-13 and SCC-39) [19]. We found enhanced TRPV1, TRPV3, ORAI1 and ORAI2 expression in SCC-13 cells (Fig. 3A). SCC39 on the other hand had detectable but reduced expression of TRPV3, ORAI1 and ORAI2. Both SCC13 and SCC-39 showed reduced expression levels of TRPC1 compared with normal keratinocytes.

**Fig. 3.**
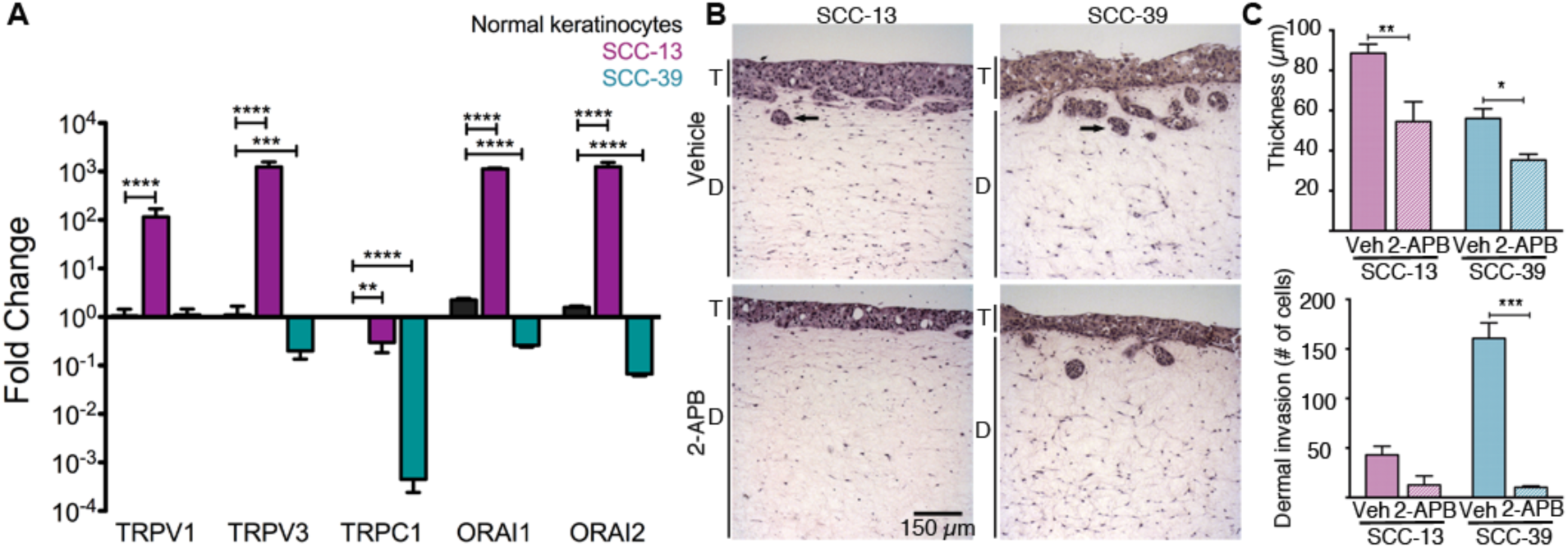
Organotypic human cSCC cultures. **A.** Transcript levels monitored by qPCR in human SCC-derived cell lines, SCC-13 (purple) and SCC-39 (blue), and normal keratinocytes (black). Fold increase compared with normal keratinocytes is plotted (N=4 technical replicates; **P<0.01, ***P<0.001, ****P<0.0001, two-way ANOVA with Bonferonni *post hoc*). Plots depict means±SDs. **B.** Organotypic human skin equivalents with SCC cell lines were treated with vehicle (top panel) or 50 µM 2-APB (bottom panel) for 14 d. **C.** SCC layer thickness (top) and number of invading cells per 10X field (bottom) were quantified for SCC-39 (blue) and SCC-13 (purple) organotypic cultures (3 rafts per treatment, T=tumor, D=dermis, *P<0.05, **P<0.01, ***P<0.0001 Bonferroni *post hoc*).

In organotypic human skin cultures, SCC-39 cells showed significantly more cells invading the dermis than SCC-13 (Fig. 3B–C; P=0.003, Student’s *t* test). SCC-39’s enhanced invasiveness is consistent with its less differentiated molecular signature [1]. Conversely, cSCC tumor formation apical to the dermis was significantly larger in SCC-13 compared with SCC-39 (Fig. 3B–C). Thus, in 3D organotypic cultures, these cSCC cell lines recapitulate a range of tumor behaviors observed in vivo.

2-APB treatment dramatically reduced tumorigenesis in human organotypic cultures seeded with each cSCC cell line. In SCC-39 organotypic cultures, 2-APB treatment inhibited cell invasion by >93% compared with vehicle-treated cultures (P<0.0001, Bonferroni *post hoc*; Fig. 3B–C). Tumor formation above the dermal equivalent in SCC-13 and SCC-39 was reduced by >37% (P<0.01, Bonferroni *post hoc*). Thus, these human preclinical models demonstrate that 2-APB reduces cSCC tumor size and dermal invasion *in vitro*.

## Discussion

Our findings identify 2-APB as a promising basis for development of targeted cSCC therapies. It is well established that calcium modulation in keratinocytes is a key step in initiating the switch from proliferation to differentiation and that this switch can be experimentally induced by increasing extracellular calcium [3, 6]. Rises in extracellular calcium are sensed through the G-protein coupled calcium-sensing receptor (CaSR), which initiates a cascade of events that mobilize intracellular calcium through both IP3-dependent store operated release and diacylglycerol-dependent activation of TRPC channels (see [6]). Intracellular calcium can also be increased by activation of TRP and ORAI channels. Here, we used a modulator of keratinocyte calcium channels to induce keratinocyte differentiation. Keratinocytes express a number of TRP and ORAI channels that have the potential to govern keratinocyte differentiation and proliferation. We found that 50 µM 2-APB caused cell-cycle arrest, a hallmark of the commitment switch from proliferation to differentiation. Moreover, even under low-calcium growth conditions, 24 h incubation with 2-APB boosted the expression of genes associated with terminal differentiation. Taken together, these findings demonstrate that 2-APB treatment is sufficient to induce early differentiation pathways in normal human keratinocytes.

Our findings extend previous genetic studies that implicate TRP and ORAI channels in epidermal differentiation [9, 10, 26, 27]. In mouse keratinocytes, TRPV3 controls late stages of keratinocyte differentiation and hair follicle morphogenesis via activation of EFGR signaling; however, a role for TRPV3 in early differentiation was previously unknown [11]. Keratinocytes also express TRPV1 and TRPC1 [14]; however, knockout animals do not demonstrate skin pathologies [28-30]. TRPC1 has been reported to promote Ca^2+^-mediated differentiation of oral epithelial cells and primary epidermal keratinocytes *in vitro* [31, 32]. ORAI1 has been shown to be essential for calcium induced differentiation [26] and mice that are mutant for ORAI1 have thick skin and hair loss [27]. Furthermore, we extend a growing body of literature that suggests ORAI channels are dysregulated in cancerous cells. ORAI1 and ORAI3 are upregulated in numerous cancers and are associated with increased proliferation, migration, and metastasis [7].

Which molecular targets mediate the effects of 2-APB on cell-cycle arrest, keratinocyte differentiation and cSCC tumor formation in vitro? Treatment with 50 µM 2-APB is well within the effective concentration ranges for the TRP channels expressed in keratinocytes (Table 1). Along with activating TRPV3, 2-APB has been reported to inhibit TRPC1; however, we do not favor a TRPC1-dependent mechanism for the proliferation arrest we observed, since a reduction in TRPC1 expression promotes keratinocyte proliferation [33]. Finally, among epidermal TRP channels, TRPV3 is the only molecular target that is not internalized or desensitized by agonist application [34]. Thus, prolonged agonist exposure is expected to constitutively increase TRPV3 channel activity levels in a dose-dependent manner over the concentration range employed. Alternatively, 2-APB modulation of TRPV3 or TRPC1 may recruit the EGFR pathway [11, 16] to govern the balance between proliferation and differentiation. Finally, 2-APB could be working through ORAI modulation to stimulate cell cycle arrest. In particular, 50 µM 2-APB inhibits ORAI1 channels (Table 1 and [35, 36]). As ORAI1 plays a key role in keratinocyte differentiation, blocking this channel with 2-APB could halt proliferation [26], [27]. Gene silencing studies are required to distinguish the effectors of 2-APB in cell-cycle arrest and induction of differentiation in human keratinocytes.

The potential role for TRP and ORAI channels in cancer has attracted much attention [37] [7] [39-41]. TRPM8 dysregulation in human prostate cancer is well established and TRPV1 knock-out mice are more susceptible to chemical skin carcinogenesis compared with wildtypes [38]. A recent study has also found that TRPV4 is downregulated in non-melanoma skin cancers [12]. ORAI1 and ORAI3 have been implicated in the progression of a number of cancers including breast cancer, melanoma, prostate cancer and glioblastoma [7]. We found that expression of TRPV3, TRPV1, TRPC1, ORAI1, ORAI2, and ORAI3 transcripts is dysregulated in human high-risk cSCC.

New therapeutic targets for high-risk cSCC are needed. Many patients treated with current standards of care suffer high rates of recurrence or disease-related deaths within five years [18]. The ability of 2-APB to arrest proliferation and promote differentiation point to its potential as a therapeutic for cSCC treatment. Although additional studies are needed to define the underlying pathophysiological mechanisms exploited by 2-APB in cSCC, our findings represent an initial step to translate their role in human keratinocyte biology into potential therapies for cSCC.

## Acknowledgements

We thank Drs. Desiree Ratner and Khan Thieu for providing SCC biopsies, Ms. Yan Lu and Ms. Rong Du for technical assistance and Dr. David Bickers for helpful discussions.

## Author contributions

Conceptualization: AMN, DMO, EAL; Methodology: DMO, EAL; Formal analysis: AMN, SAG, YM; Investigation: AMN, SAG, MER; Resources: UBJ, DMO; Writing – original draft: AMN, YM; Writing – review & editing: AMN, SAG, YM, MER, UBJ, DMO, EAL; Visualization: AMN, YM; Supervision: EAL; Project Administration: EAL; Funding acquisition: DMO, EAL.

